# Mast Cell Derived Histamine Negatively Regulates Hematopoiesis

**DOI:** 10.1101/2022.02.03.479012

**Authors:** Bailey R. Klein, Julianne N.P. Smith, Ramachandra Katabathula, Rahul Chaudhary, Zhenxiang Gao, Brittany A. Cordova, Frederick Petroze, Riya Tiwari, Stanton L. Gerson, Rong Xu, Sanford D. Markowitz, Amar B. Desai

**Author notes:** Correspondence: Amar B. Desai, Department of Medicine, Case Western Reserve University. Author Contribution: BRK conceived and designed the study, collected and interpreted the data, and wrote the manuscript. JNPS conceived and designed the study and collected and interpreted the data. RK designed studies, interpreted data, and contributed to manuscript writing. RC collected the data. ZG collected and interpreted the data. BAC collected the data. FP collected the data. RT collected the data. SLG designed studies and interpreted data. RX designed studies, interpreted data, and contributed to manuscript writing. SDM conceived and designed studies and interpreted data. ABD conceived and designed the study, interpreted the data, wrote the manuscript, and provided final approval of the manuscript.

## Abstract

Hematopoietic stem cells (HSCs) are essential for generating all blood cell types and maintaining immune function and oxygen transport. This requires tight regulation of self-renewal, differentiation, and quiescence, driven by intrinsic and extrinsic signals. While the influence of many HSC progeny on HSC decisions are recognized, the role of mast cells (MCs) remain understudied. MCs are known for their immunomodulatory functions through the secretion of factors such as histamine and could offer new insights into HSC regulation. In this study, we describe a novel role for MC-derived histamine in modulating HSC behavior. We observed that genetically MC-deficient “SASH” mice exhibit increased hematopoietic output and bone marrow (BM) HSCs, characterized by an enhanced quiescent signature that increases resistance to myeloablative chemotherapy. The SASH microenvironment also contained increased frequencies of HSC-supportive cell types and expression of genes conducive to HSC maintenance, which together accelerated HSC engraftment when wild-type BM was transplanted into SASH recipients. Moreover, we found lower serum histamine levels in SASH mice, and that the enhanced hematopoietic phenotype observed in these mice could be reversed by administering exogenous histamine. Subsequent experiments with FDA-approved antihistamines in wild-type mice revealed that cetirizine, an H1R inverse agonist, notably increased HSC frequency in the BM. Overall, our findings implicate MCs are negative regulators of HSC function. This lays the groundwork for future studies to elucidate the underlying mechanisms and explore the therapeutic potential of modulating histamine signaling to promote hematopoiesis.

## INTRODUCTION

Hematopoietic stem cells (HSCs) are primitive multipotent cells that reside in the bone marrow (BM) and differentiate into all the cellular components of blood, including myeloid, megakaryocytic, lymphoid, and erythroid lineage cells^1^. This process of differentiation, known as hematopoiesis, occurs life-long from a small pool of self-renewing HSCs, which employ protective mechanisms to ensure lifelong functionality. These include residing in a hypoxic niche to minimize exposure to reactive oxygen species, maintaining a quiescent state to protect themselves from genetic insults, and the undergoing apoptosis in response to excessive cellular damage^2–4^. The BM stroma consists of mesenchymal cells, endothelial cells, and some HSC progeny, such as megakaryocytes and macrophages. It provides a supportive microenvironment for HSC functions by producing essential factors including stem cell factor (SCF), CXCL12, and thrombopoietin (TPO), which are necessary to maintain the BM niche^5–9^. Among these cellular interactions and signaling mechanisms, recent discoveries have highlighted a distinct role for histamine derived from bone marrow myeloid cells. Specifically, histamine has been found to induce quiescence and self-renewal by activating the H2 receptor on myeloid-biased HSCs, pointing to a nuanced interplay of local and systemic factors that collectively safeguard HSC integrity^10^. Currently, there are no studies linking mast cells (MCs), which are one of the largest producers of histamine, to HSC function. This may be due to a lack of proximity as MC precursors originate in the BM, but then migrate to peripheral tissues for further development, resulting in few to no mature MCs within the BM^11–14^.

MCs are tissue-resident immune cells of the myeloid lineage located in mucosal and epithelial tissues throughout the body^15^. These cells are canonically known to be first responders to infection and mediators of allergy through activation of FcεRI and pattern recognition receptors^16, 17^. Activation of MCs causes a release of mediators such as proteases, leukotrienes, and histamine that elicit physiological responses including vasodilation, inflammatory cell recruitment, and angiogenesis^18, 19^. Degranulation releases copious amounts of histamine, but MCs also constitutively secrete a basal level of histamine systemically where it can bind and activate one of its four distinct cognate G-coupled protein receptors: H1R, H2R, H3R, and H4R^20–23^. Antihistamines, a well-tolerated class of drugs, bind to these receptors to inhibit their activation and physiological effects. H1R and H2R antihistamines, approved by the FDA and used by millions of people daily are used to treat allergy symptoms and gastric ulcers respectively^24, 25^.

In this study, we showcase a unique phenotype wherein MCs exhibit a negative regulatory influence on hematopoiesis and hematopoietic output. Additionally, we present evidence that this phenotype can be replicated pharmacologically by targeting H1R with the FDA approved antihistamine cetirizine. These findings suggest a potential strategy for enhancing hematopoietic output in multiple scenarios including chemotherapy induced myelosuppression, transplantation, and bone marrow failure.

## RESULTS

### MC deficient Kit^W-sh/W-sh^ “SASH” mice display leukocytosis and BM HSPC expansion

We characterized the steady state hematopoietic compartments of the commercially available MC deficient mouse model (SASH c-Kit^w-sh^, Jackson Labs Stock #030764). The Kit^W-sh/W-sh^ (hereinafter referred to as SASH) mice harbor an inversion mutation impacting the transcriptional regulatory elements of the *c-Kit* gene leading to a total ablation of their MC population shortly after birth^26, 27^. Complete blood count analysis on peripheral blood demonstrated SASH mice have a 1.2-fold increase in total white blood cells (WBCs), 1.8-fold increase in neutrophils (NE), 1.6-fold increase in monocytes (MO), and 1.4-fold increase in platelets (PLT) (Figure 1A). Despite the increases in peripheral blood WBCs and specific myeloid populations, we found no impact of MC deficiency on peripheral blood lymphocytes (LY) or RBCs (Supplemental Figure 1). In the BM, we observed a 1.4-fold increase in total cellularity and through immunophenotypic analysis (gating strategy outlined in Supplemental Figure 7) we identified significant 1.7-fold increase in lineage^-^Sca-1^+^c-Kit^+^ (LSK) cells and 2.1-fold increase in CD150^+^CD48^-^ long-term HSCs (LT-HSC) (Figure 1B). Additionally, we investigated the myeloid and lymphoid frequencies within the BM (gating strategy outlined in Supplemental Figure 8) and observed a significant increase in total myeloid frequency and a significant decrease in B cells and CD8^+^T cells (Supplemental Figure 1).

**Figure 1:**
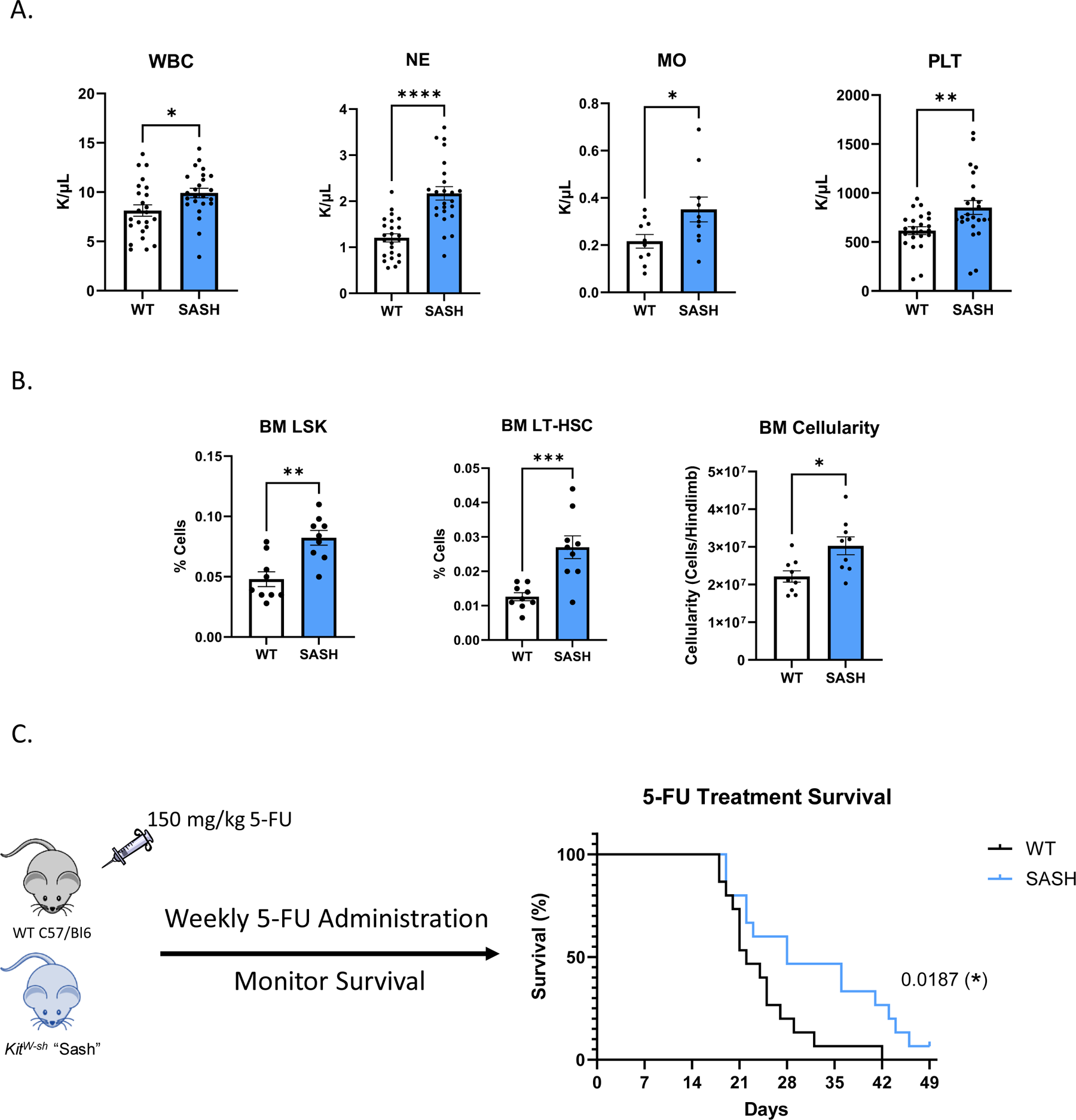
MC deficient KitW-sh/W-sh “SASH” mice display leukocytosis and BM HSPC expansion. **A**. Complete blood count analysis of circulating WBC, MO, NE, and PLT at steady-state in WT and SASH mice. N=24 mice/group, except monocytes was N=10 mice/group. **B.** Immunophenotypic analysis of HSPCs and LSKs per hindlimb. N=9 mice/arm. Total nucleated bone marrow cells per hindlimb were obtained. N=9 mice/group. **C.** Survival curve of mice subjected to weekly 150 mg/kg injections of 5-FU. N=15 mice/group. **A, B.** Error bars represent SEM. Students two-tailed T test was used for statistical analysis with p>0.05=*, p>0.01=**, p>0.001=***,p>0.0001=****

We additionally characterized spleens of SASH mice and found these mice displayed splenic extramedullary hematopoiesis as indicated by significant increases in splenic HSPCs and CFU potential (Supplemental Figure 2). This is consistent with previously reported findings of extramedullary hematopoiesis in the spleen of SASH mice^28^. Taken together, these data demonstrate significantly enhanced hematopoietic output following loss of MCs. To next test the functional consequence of the observed increase in hematopoietic output we assessed the capacity of SASH mice to recover following repeated 5-Fluorouracil (5-FU) administration. 5-FU, a pyrimidine analog, disrupts DNA synthesis and repair, preferentially targeting and eliminating actively cycling cells within the hematopoietic system. Therefore, 5-FU can be utilized in hematopoietic recovery studies to induce genotoxic stress, creating a selective pressure against cycling HSPCs. Consequently, the ability to recover from 5-FU relies on the reserve of quiescent HSCs to replenish the depleted hematopoietic cell populations. This makes 5-FU an ideal agent for evaluating the depth of the HSC pool, as resistance to its cytotoxic effects suggests the presence of a substantial reservoir of quiescent HSCs^29, 30^. Indeed, we observed that genetic loss of MCs resulted in a significant survival extension compared to WT mice following weekly 150 mg/kg 5-FU administrations (Figure 1C), indicating that the increased phenotypic HSCs and progenitors are functional and confer protection against hematopoietic injury.

### SASH lineage depleted BM exhibit transcriptional signatures indicative of HSPC quiescence

Next, to elucidate the observed hematopoietic phenotype in SASH mice we performed RNA sequencing on both bulk BM cells and lineage depleted BM cells (a population enriched for hematopoietic stem and progenitor cells) from both WT and SASH mice (Figure 2A). Principal component analysis plots indicate no obvious stratification between bulk bone marrow populations of SASH and WT mice (teal vs. red). However, lineage depleted populations demonstrated a clear divergence in expression profiles (green vs. purple), suggesting a unique HSPCs signature in SASH mice (Figure 2B). Gene set enrichment analyses of lineage depleted cells (Supplemental Figure 3) in SASH mice revealed a downregulation in pathways related to cellular proliferation and division, such as “DNA replication”, “cell cycle checkpoint signaling”, and “nuclear chromosome segregation” coupled with an increase in “negative regulation of cell activation” within the M5 ontology gene set (BP) defined by the Gene Ontology consortium (Figure 2C). This signature is consistent with our 5-FU data, demonstrating a quiescent transcriptional signature within the lineage depleted portion of the BM. Moreover, the observed upregulation in “leukocyte migration” and “myeloid leukocyte activation” pathways indicates that, alongside a quiescent transcriptional profile, these cells are primed for WBC production (Figure 2D). This suggests a balance between maintaining quiescence and the potential for activation.

**Figure 2:**
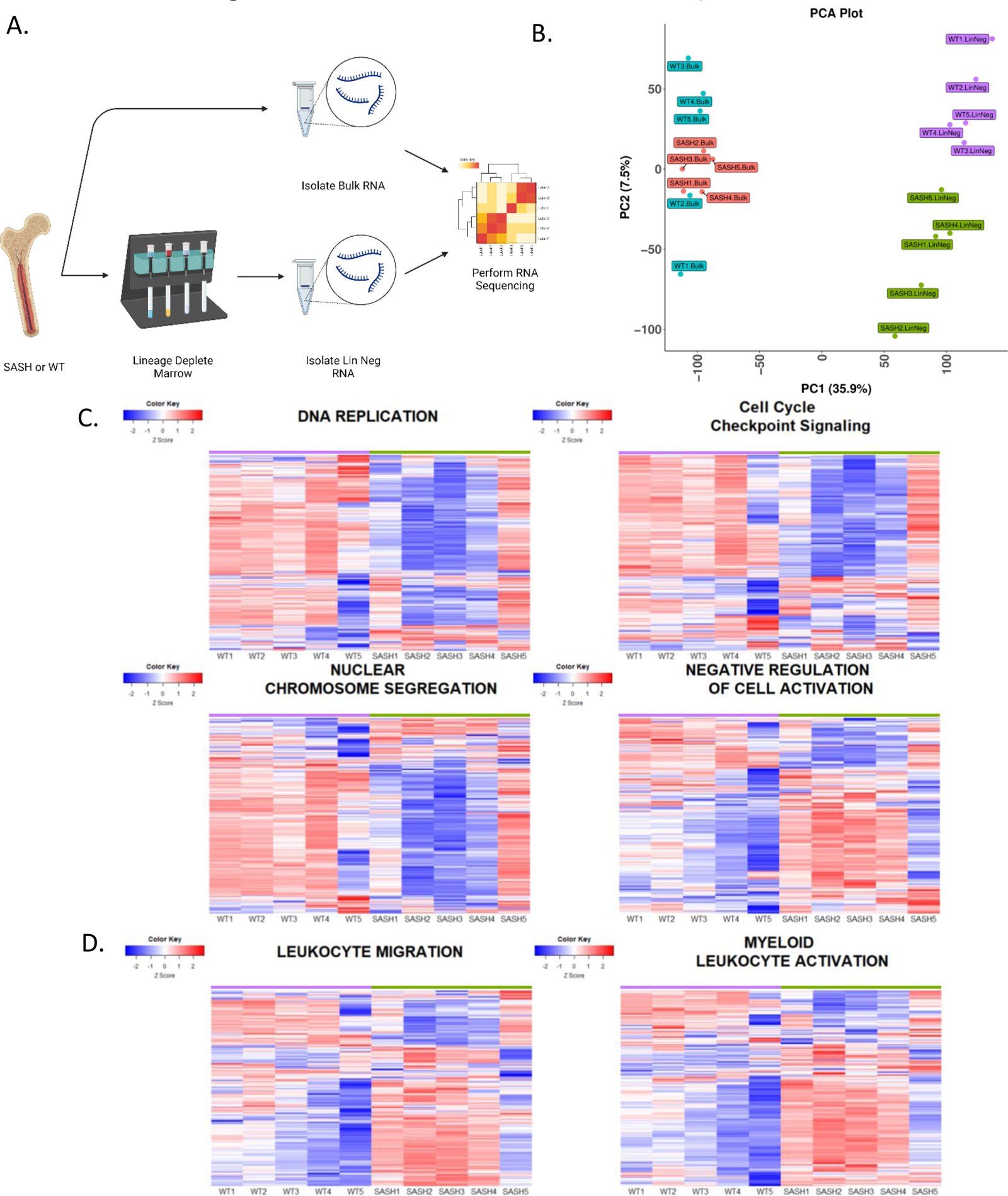
SASH lineage depleted BM exhibit transcriptional signatures indicative of HSPC quiescence. **A.** Schematic depicting the experimental design for RNA sequencing study. **B.** Principal component analysis (PCA) plot of SASH and WT bulk and lineage depleted populations. **C,D.** Heatmap of top differentially regulated genes between SASH and WT mice showing **C.** pathways related to cellular activation and replication **D.** pathways related to WBC activation and migration. N=5 mice/group.

### The SASH BM microenvironment is primed for hematopoietic engraftment

We next investigated the frequency of HSC supportive cell types through immunophenotypic characterization within the CD45^-^/Ter119^-^ BM population (gating strategy outlined in Supplemental Figure 9). We found that leptin receptor positive cells (LepR^+^), had a 1.4-fold increase and adipocytes had a 1.9-fold increase in frequency within SASH BM (Figure 3A). LepR+ cells, which comprise skeletal stem cells, along with osteogenic and adipogenic progenitor cells are pivotal for their dual role in the niche; they not only contribute to the physical scaffolding that determines the structural integrity of the niche but also secrete a range of soluble factors that are vital for HSC maintenance, such as cytokines and growth factors. Similarly, the increase in adipocytes, which are known to secrete adipokines and other regulatory molecules, suggests an enriched environment for HSC support^31^.

**Figure 3:**
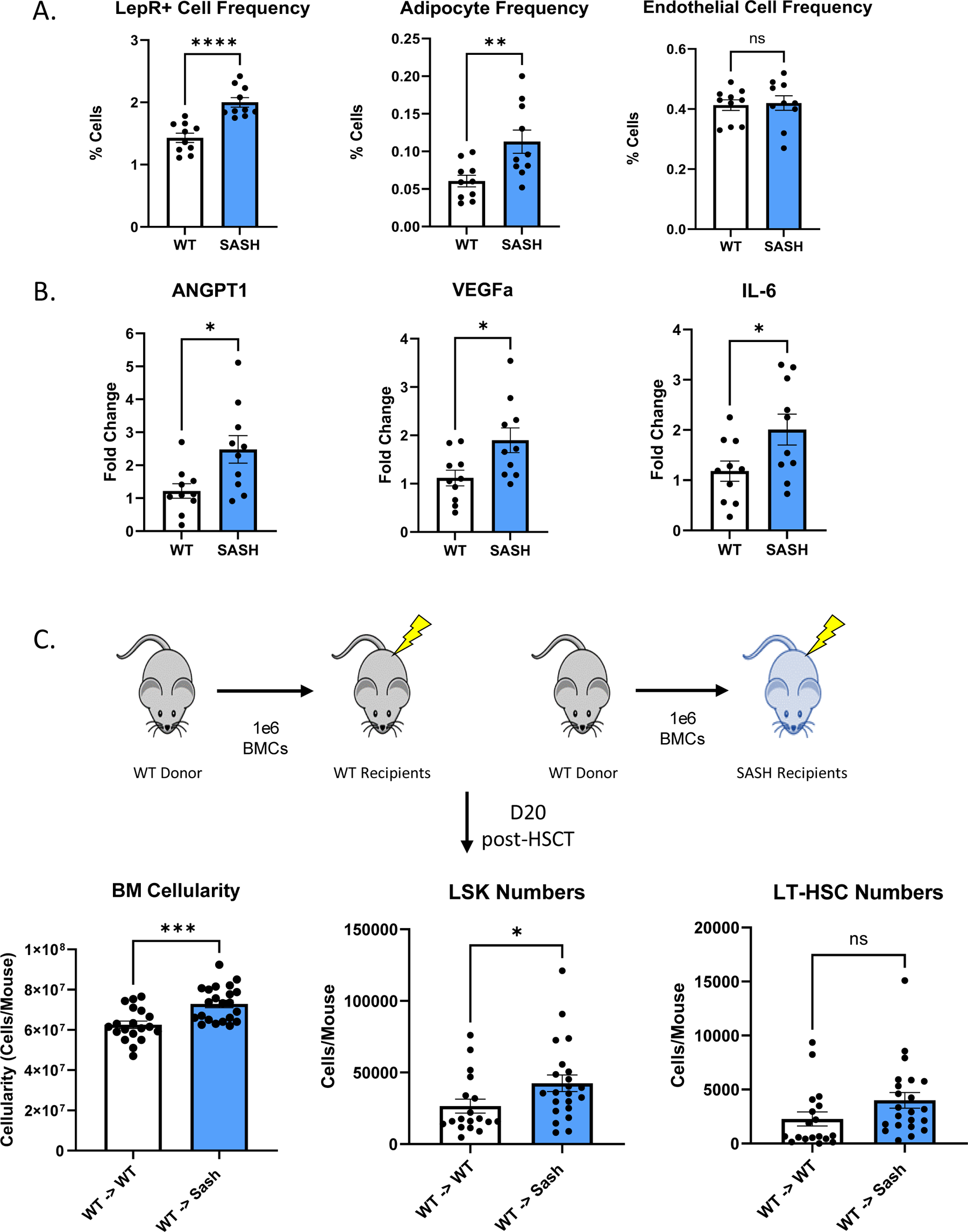
The SASH BM microenvironment is primed for hematopoietic engraftment. **A.** Flow cytometric analysis of BM stromal populations. N=10 mice/arm. **B.** RT-PCR analysis of niche factor genes within BM stromal cells. N=10 mice/arm. **C.** BM cellularity and quantification of LSK and LT-HSC cells per mouse (2 hindlimbs) of recipients 20 days post-transplant. N=18-22 mice/arm. **A-C.** Error bars represent SEM. Students two-tailed T test was used for statistical analysis with p>0.05=*, p>0.01=**, p>0.001=***,p>0.0001=****

We next investigated a panel of different niche genes within the CD45^-^ population of the BM (Supplemental Figure 4) and observed significant increases in angiopoietin-1 (*Angpt1*) (important for maintaining a quiescent state within HSCs^32^, and notably secreted by LepR+ cells^33^), *Vegfa* (important for HSC survival^34^), and *Il6* (needed for granulopoiesis^35^) (Figure 3B). To determine the functional effects of the observed gene expression data, we performed transplantation studies in which WT donor BM was transplanted either into WT or SASH recipient mice. We sacrificed mice on day 20 following transplantation and found 1.2-fold higher BM cellularity in SASH recipients compared to WT controls, 1.6-fold greater LSKs and trends towards increased total LT-HSC (1.76-fold) (Figure 3C). This data suggests that even after myeloablation, the SASH microenvironment is more favorable to hematopoietic engraftment and thus peripheral reconstitution following transplantation compared to WT counterparts.

### SASH mice contain lower histamine levels and histamine administration impairs hematopoietic output

To determine the role of histamine in modulating hematopoietic output in SASH mice we compared the steady-state levels of serum histamine between SASH and WT mice and observed that WT mice contained 1.6-fold higher histamine than SASH (Figure 4A). To further investigate this connection, we assessed the expression of histidine decarboxylase (*Hdc*), the sole enzyme that converts L-histidine to histamine, thus serving as a specific marker for the biosynthesis of histamine^36^. We observed no significant differences in *Hdc* expression between SASH and WT mice in both bulk BM and CD45^-^ populations, suggesting that the decrease in histamine levels was due specifically to the loss of MCs (Figure 4B). Moreover, we investigated the expression levels of *Hrh1*, *Hrh2*, and *Cysltr1* in both bulk and CD45^-^ populations in SASH and WT mice (Supplemental Figure 5). Interestingly, we observed a decrease in the expression of *Hrh1* within the CD45^-^ population of SASH BM, indicating reduced H1-mediated signaling in response to histamine in these cells.

**Figure 4:**
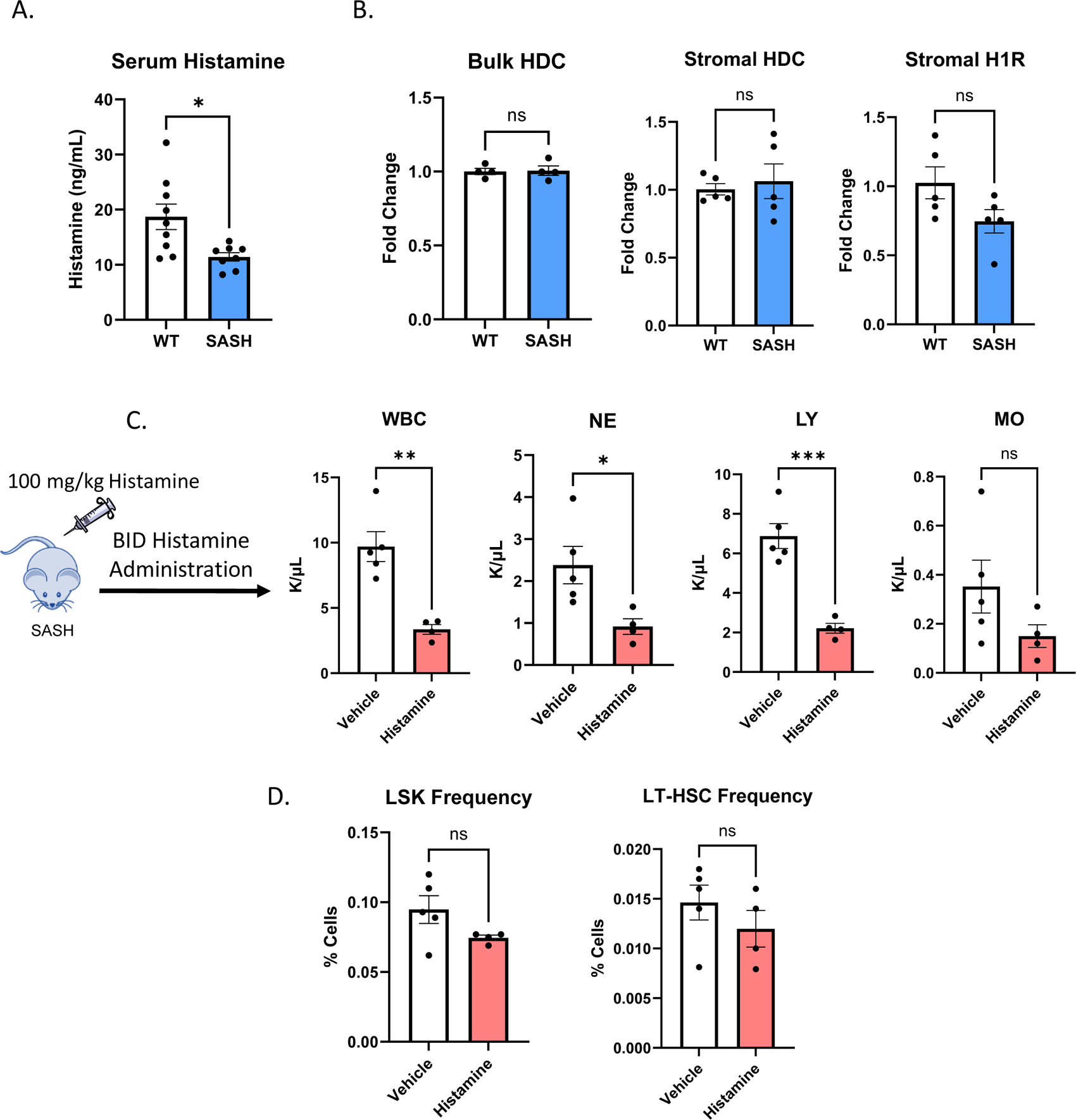
SASH mice contain lower histamine levels and histamine administration impairs hematopoietic output. **A.** ELISA analysis of histamine concertation in the serum of WT and SASH mice. N=8-9 mice/arm. **B.** RT-PCR analysis of HDC and H1R within bulk and stromal BM cells. N=4-5 mice/arm. **C.** Complete blood count analysis of circulating WBC, NE, LY, and MO after 7 days of BID 100 mpk histamine administration. N=4-5 mice/group. **D.** Immunophenotypic analysis of HSPCs and LSKs per hindlimb after 7 days of BID 100 mpk histamine administration. N=4-5 mice/arm. **A-D.** Error bars represent SEM. Students two-tailed T test was used for statistical analysis with p>0.05=*, p>0.01=**, p>0.001=***,p>0.0001=****

To explore the functional consequences of reduced histamine in SASH mice, we administered exogenous histamine to these animals and evaluated its effects on peripheral blood counts and BM HSPC frequencies. Treatment of SASH mice at 100 mg/kg histamine BID for 7 days resulted in a 2.9, 2.6, and 3.1-fold reduction in WBC, NE, and LY respectively, demonstrating that histamine supplementation reduces peripheral blood output in SASH mice (Figure 4C). Notably, RBC and PLT counts were not altered by the 7-day histamine treatment (Supplemental Figure 6). We tested a lower dose of 10 mg/kg and a 4-day blood analysis time point on both doses but saw no alterations (Supplemental Figure 6). We also examined BM HSPCs and observed a trend towards decreased cell populations only at the higher histamine dose (Figure 4D and supplemental Figure 6). Together this data suggests histamine negatively regulates hematopoietic function and the loss of MC derived histamine in SASH mice contributes to the observed increase in hematopoietic output.

### *In vivo* antihistamine treatment results in steady-state BM progenitor cell expansion

Having identified a potential causative role for histamine in driving the hematopoietic phenotype in SASH mice, we sought to pharmacologically mimic this effect using antihistamines. Mice were treated twice daily for 7 days with a selection of FDA-approved antihistamines. Specifically, we administered 20 mg/kg of cetirizine (H1R inverse agonist^37^), 10 mg/kg of famotidine (H2R antagonist^38^), 5 mg/kg of montelukast (leukotriene receptor antagonist^39^), and 10 mg/kg of ketotifen (H1R inverse agonist and MC stabilizer which prevents the release of additional inflammatory mediators from MCs^40^). While additional studies will be required to identify optimal dosing regimens, our initial findings revealed a statistically significant 1.7-fold increase in both LSK and LT-HSC populations specifically following cetirizine treatment (Figure 5B). In addition, we performed a myeloid and lymphoid population analysis within the BM and found that cetirizine induced significant changes including an increase in total myeloid cells, B cells, and CD8^+^ T cells (Figure 5C).

**Figure 5:**
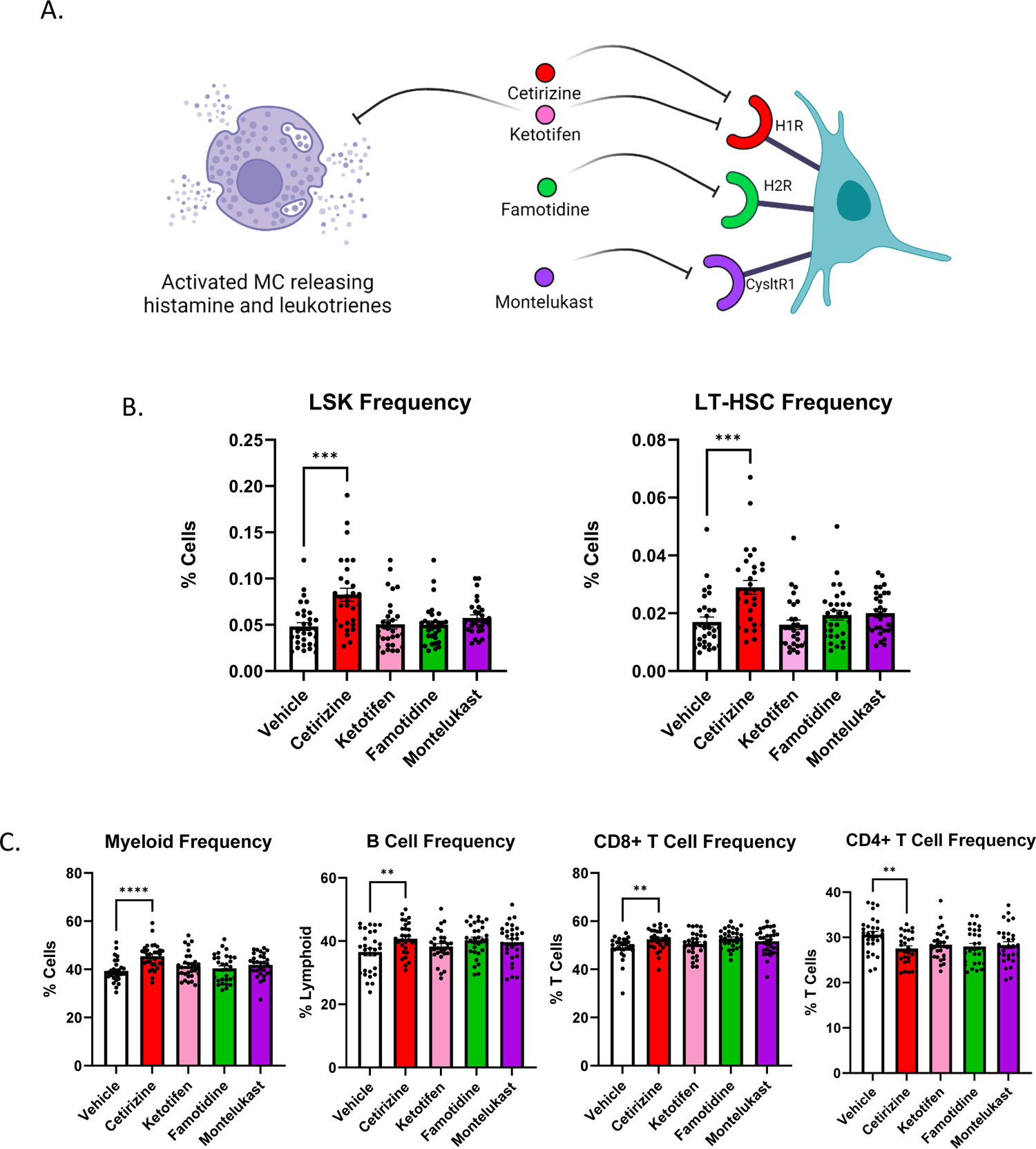
*In vivo* antihistamine treatment results in BM progenitor cell expansion at steady-state. **A**. Schematic depicting the mechanism of action for FDA-approved drugs that target histamine and leukotriene receptors and for drugs that stabilize MCs. **B.** Immunophenotypic analysis of HSPCs and LSKs per hindlimb of WT mice treated BID with 2, 1, 0.5, and 1 mg/kg of cetirizine, famotidine, montelukast, and ketotifen respectively. N=30 mice/arm. **C.** Immunophenotypic analysis of myeloid, B and T cells per hindlimb of the same mice in B. N=30 mice/arm (15/sex). **B, C.** Error bars represent SEM. Students two-tailed T test was used for statistical analysis with p>0.05=*, p>0.01=**, p>0.001=***,p>0.0001=****

### EHR data identifies association between antihistamine use and white blood cell counts

To determine the clinical relevance of our murine data, we analyzed electronic health record (EHR) data from TrinetX in patients diagnosed with allergy and prescribed antihistamines vs other anti-allergy medications. The ‘exposure risk’ represents the proportion of patients taking antihistamines who had elevated white blood cell counts. Conversely, the ‘control risk’ refers to the proportion of patients taking other anti-allergy medications who had elevated WBC counts. Antihistamines were associated with a significantly increased risk for elevated white blood cell counts compared with other anti-allergy medications during a 1-year follow-up in patients with allergy and no history of elevated white blood cell counts (HR: 2.09, 95% CI: 1.57-2.78), suggesting a strong association in a relative short time follow-up (**Table 1**). While this data does not establish causality, it intriguingly parallels our animal model results and prompts further investigation into whether this association is indicative of a conserved biological mechanism. It also provides a compelling link between antihistamine use and hematopoietic output in a clinical setting, thereby reinforcing the translational relevance of our animal model findings.

**Table 1.** Outcome: elevated white blood cell count (follow up: 1 year)

| *Table 1. Outcome: elevated white blood cell count (follow up: 1 year)* |  |  |  |
| --- | --- | --- | --- |
| Cohort setting | Exposure risk | Control Risk | Hazard Ratio (95% CI) |
| Antihistamines group vs<br>Decongestants and<br>antiallergics group | 1.15% | 0.55% | 2.09 (1.23,3.56) |
| Antihistamines group vs<br>Corticosteroids group | 0.58% | 0.27% | 2.09 (1.57,2.78) |

## DISCUSSION

This work elucidates a previously unexplored regulatory mechanism within the hematopoietic system demonstrating how MCs and histamine signaling via H1R influence both HSC function and overall hematopoietic output. We found that MC deficient SASH mice display leukocytosis, expanded HSPC populations in the BM, and increased splenic extramedullary hematopoiesis. Our transcriptomic analysis indicated that the lineage depleted BM is predisposed to increased WBC output without compromising its quiescent state. Increased resistance to repeated 5-FU treatment validated this predisposition. Further analyses of cell frequencies and gene signatures within the BM niche suggested a more supportive microenvironment, which correlated with improved engraftment following transplantation into SASH recipient animals.

Our work additionally posits that the reduced histamine levels observed in SASH mice contribute to their hematopoietic phenotype. We demonstrate that administration of exogenous histamine reverses the expanded peripheral blood output and BM HSPCs. Collectively these findings indicate that histamine acts a negative regulator of hematopoietic function.

In addition to elucidating a novel mechanism in SASH mice by which MC derived histamine regulates hematopoiesis, we also found that inhibition of histamine signaling with cetirizine treatment, an H1R inverse agonist, resulted in significant expansion of HSPC populations in the BM, alongside increased myeloid populations and lymphoid cell subsets. Importantly we also identified a consistent result from retrospective analysis of electronic health record data, in which patients taking antihistamines had significantly higher white blood cell counts than untreated cohorts.

Our data suggests cetirizine could be utilized as a novel approach to promote hematopoietic function. Repurposing FDA-approved therapies for alternative disease indications offers advantages such as shorter development time (5-7 years), a higher rate of approval for the alternative indication, a substantial reduction in development costs (ranging from 40-70%), and access to a wealth of existing data from the extensive FDA approval process these compounds have undergone^41^. Based on our findings, this work may have implications for several clinical needs including:

- Poor graft function (PGF)-a complication of allogeneic transplantation characterized by unexplained persistent cytopenia despite full donor chimerism which occurs in 5 to 27 percent of patients and is associated with increased morbidity and mortality^42^. The management of PGF is limited and often ineffective, typically involving supportive treatments such as hematopoietic growth factors (like G-CSF), eltrombopag, and stem cell reinfusion or boosts. By bolstering the HSC pool, cetirizine could effectively augment hematopoietic regeneration, potentially mitigating the persistent cytopenia characteristic of PGF and improving overall patient outcomes post-transplant.
- Myelofibrosis-a chronic myeloproliferative disorder marked by progressive scarring of the bone marrow, leading to severe anemia, weakness, fatigue, and often, splenomeglia^43^. MF patients also display clonal hematopoiesis, which is characterized by HSC production of a small population of dominant clones rather than a healthy polyclonal population, and which is associated with a significantly higher risk of transformation^44^. While targeting MCs in fibrotic disease has recently been proposed as a novel therapeutic strategy^45, 46^, there is a possibility that antihistamines may display dual efficacy in MF by acting to not only limit clonal dominance but also reduce fibrotic onset.

Overall, this work uncovers a previously unrecognized role of MCs in regulating hematopoiesis and provides insights into the underlying molecular mechanisms. Further investigations into the precise mechanisms mediating MC-HSPC interactions are warranted, along with exploring the therapeutic potential of MC-targeted interventions in additional settings of insufficient hematopoiesis including chemotherapy induced neutropenia, various anemias, and transplantation settings. This research not only contributes to a deeper understanding of the mechanisms governing normal and regenerative hematopoiesis, but also demonstrates the potential of repurposing MC targeting therapies as a cost effective method to promote hematopoietic function in the clinical setting.

## METHODS

### Animals

Animals were housed in the AAALAC-accredited facilities of the CWRU School of Medicine. The Case Western Reserve University Institutional Animal Care and Use Committee (IACUC) approved the husbandry and experimental procedures in accordance with approved IACUC protocols 2019-0065. C-Kit^W-sh^ mice were purchased from The Jackson Laboratory. Littermate wild-type control animals were used as comparator animals for all studies, and a combination of male and female mice, aged 8-12 weeks old, were used for all studies. Mice were housed in standard microisolator cages and maintained on a defined, irradiated diet and autoclaved water. All animals were observed daily for signs of illness. Animals that were used for tissue analysis were sacrificed humanely with CO_2_ and then by cervical dislocation. Bone marrow was obtained by removing the tibia and femur and flushing the cells into FACS buffer. Splenocytes were obtained by removing the spleen and mincing them into a single-cell suspension.

### Complete Blood Count Analysis

Peripheral blood was collected by submandibular bleeding into Microtainer EDTA tubes (Becton-Dickson cat # 365974). Blood counts were analyzed by a Hemavet 950 FS hematology analyzer.

### Flow Cytometry

Cells were resuspended and stained with 200 µL of FACS with fluorescently labeled antibodies for 15 minutes and then fixed in 1% PFA for 15 minutes. Samples were then washed with FACS buffer and finally resuspended in 200 µL of FACS buffer. Data was acquired on an LSRII flow cytometer (BD Biosciences). Analysis was performed on FlowJo software (TreeStar).

### 5-FU Administration

C-Kit^W-sh^ and WT mice were treated with 150 mg/kg of 5-FU (Sigma cat # 7F6627) in a 10% DMSO solution in PBS every seven days via IP injection. Mice were monitored daily and sacrificed when they exhibited >25% weight loss or became moribund. Animal survival was monitored throughout this study.

### Histamine ELISAs

Peripheral blood was collected from mice and allowed to clot before centrifugation to collect serum. Samples were frozen until day of assay. ELISA assays performed using Abcam kits (cat # AB213975) according to manufacturer’s protocol.

### Histamine and Antihistamine Administration

Histamine dihydrochloride was solubilized directly into PBS. Cetirizine dihydrochloride (Sigma cat # C3618), ketotifen fumarate salt (Sigma cat # K2628), famotidine (Sigma cat # F6889), and montelukast sodium (Sigma cat # PHR1603) were dissolved in DMSO at 60, 30, 30 and 15 mg/mL respectively. Each compound was then diluted in water to a final concentration of 2, 1, 1, and 0.5 mg/mL respectively, diluting the DMSO to 3.3%. Naïve SASH C57BL/6 were administered histamine and antihistamines respectively via IP injection on a BID schedule (A.M. and P.M. administration) for 7 days and were sacrificed 1 hour after the 15^th^ dose.

### Bone Marrow Transplantation

Mice were exposed to a single dose of 10 Gy total body irradiation from a cesium source. 16-24 hrs later, mice received 1e6 whole bone marrow cells by retroorbital injection. Recipients were monitored daily for signs of distress or illness and sacrificed on day 20 to assess bone marrow and spleen cellularity in addition to cytometric assessment of cell populations.

## Acknowledgments

This work was supported by NIH grants R00 HL135740, RM1GM42002, R21AG075573, and by the Radiation Resources Core Facility, the Translational Therapeutics Core Facility, the Hematopoietic Biorepository and Cellular Therapy Core Facility, and the Cytometry & Microscopy Core Facility of the Case Comprehensive Cancer Center (P30CA043703). This work was also supported by a generous award from the Ohio Cancer Research Foundation. This project was also supported by the Clinical and Translational Science Collaborative of Northern Ohio, which is funded by the National Institutes of Health, National Center for Advancing Translational Sciences, Clinical and Translational Science Award grant, UM1TR004528.

