## Supplemental Materials for "Mast Cell Derived Histamine Negatively Regulates Hematopoiesis"

### Supplemental Figure 1: Genetic loss of MCs does not impact PB leukocytes or RBC counts but does impact BM myeloid and lymphoid populations

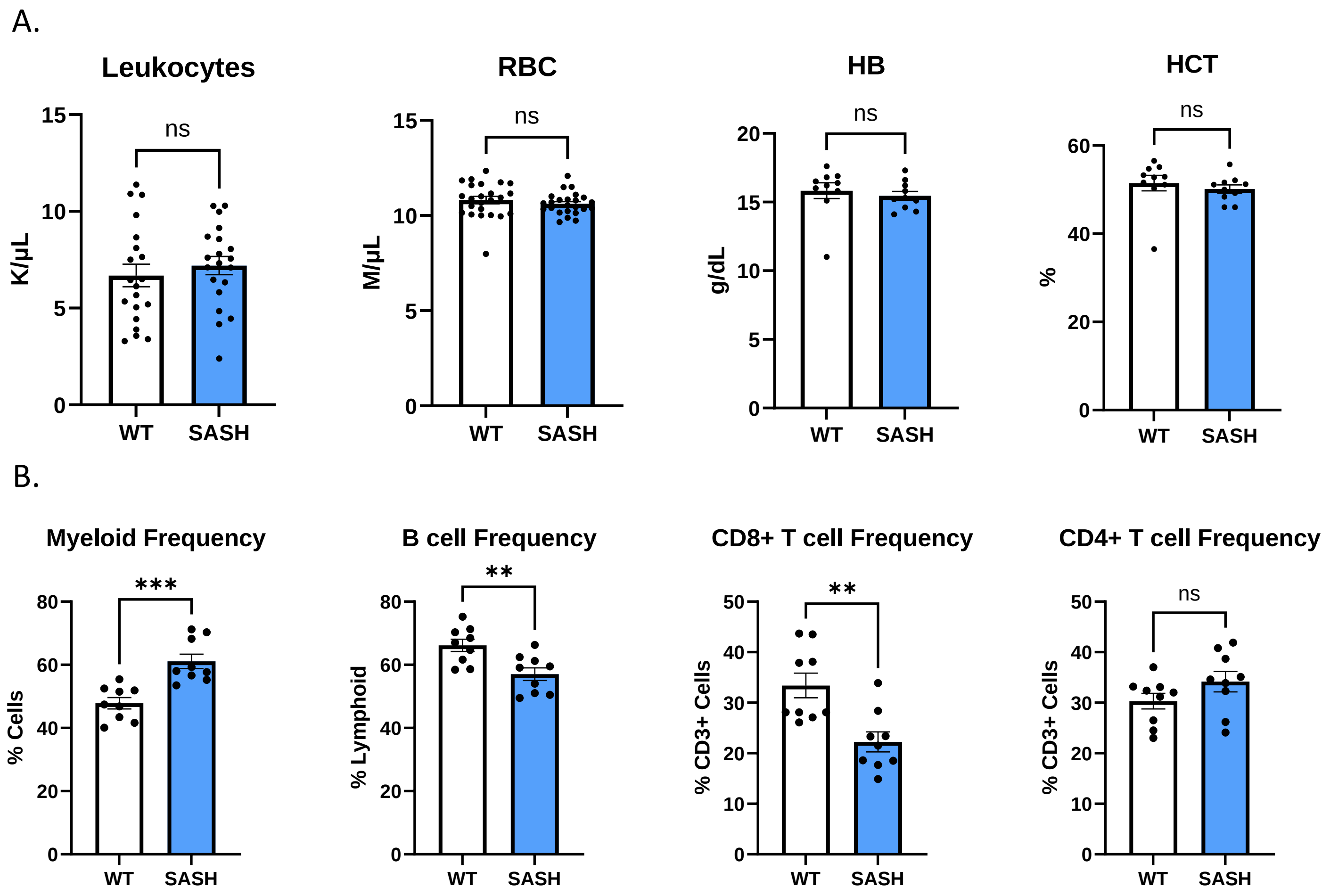


**Supplementary Figure 1: A**. Complete blood count analysis of circulating LY, RBC, Hb, and HCT at steady-state in WT and SASH mice. N=10-24 mice/group. **B.** Immunophenotypic analysis of myeloid and lymphoid populations per hindlimb. N=9 mice/arm. **A, B.** Error bars represent SEM. Students two-tailed T test was used for statistical analysis with p>0.05=*, p>0.01=**, p>0.001=***,p>0.0001=****

### Supplemental Figure 2: SASH spleens display increased extramedullary hematopoiesis

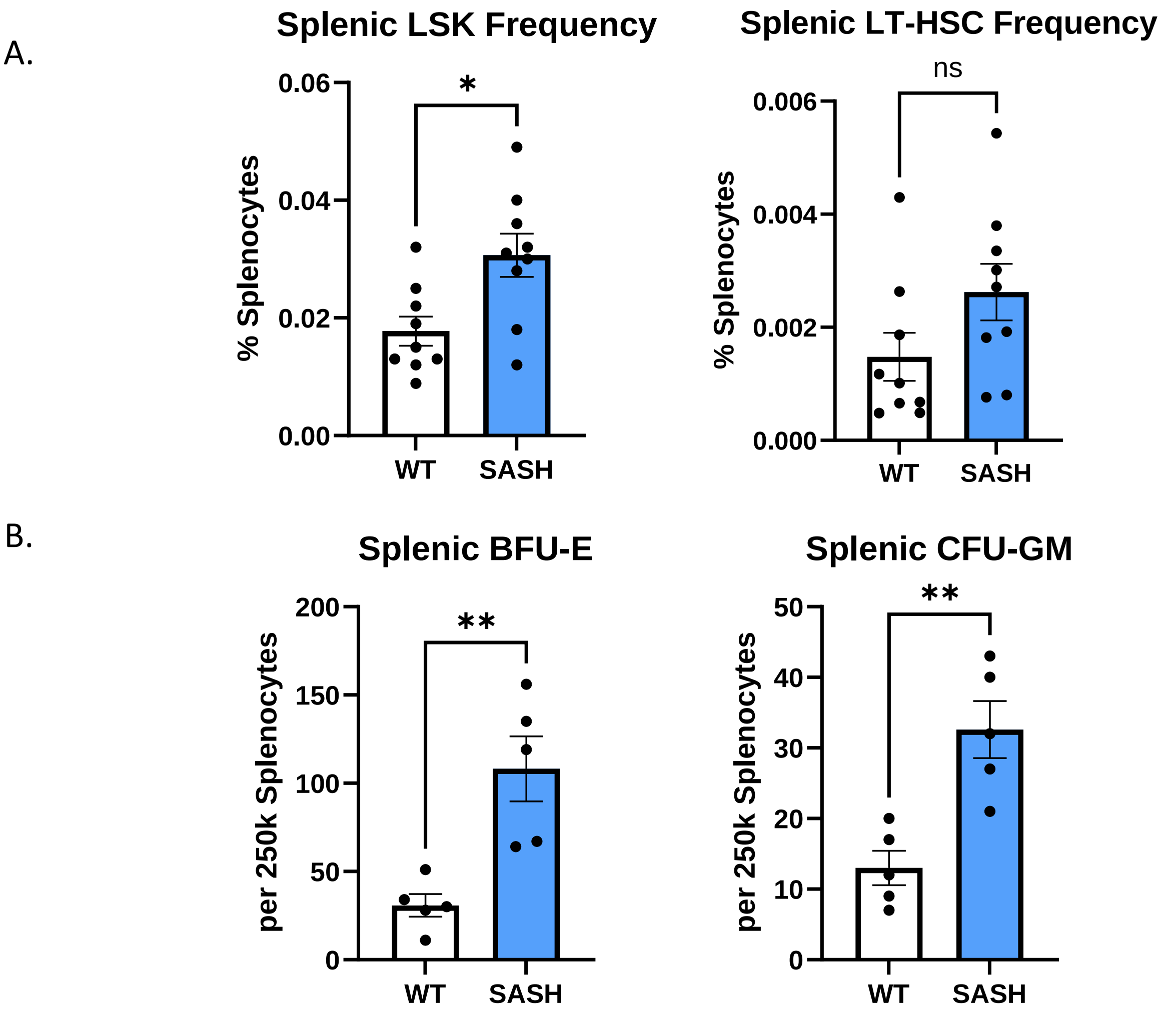


**Supplementary Figure 2: A**. Immunophenotypic analysis of LSKs and HSPCs per spleen. N=9 mice/arm. **B.** Methylcellulose assay CFU output from 250,000 splenocytes . N=5 mice/arm. **A, B.** Error bars represent SEM. Students two-tailed T test was used for statistical analysis with p>0.05=*, p>0.01=**, p>0.001=***,p>0.0001=****

### Supplemental Figure 3: SASH mice differentially express pathways related to cell proliferation, migration, and immunity

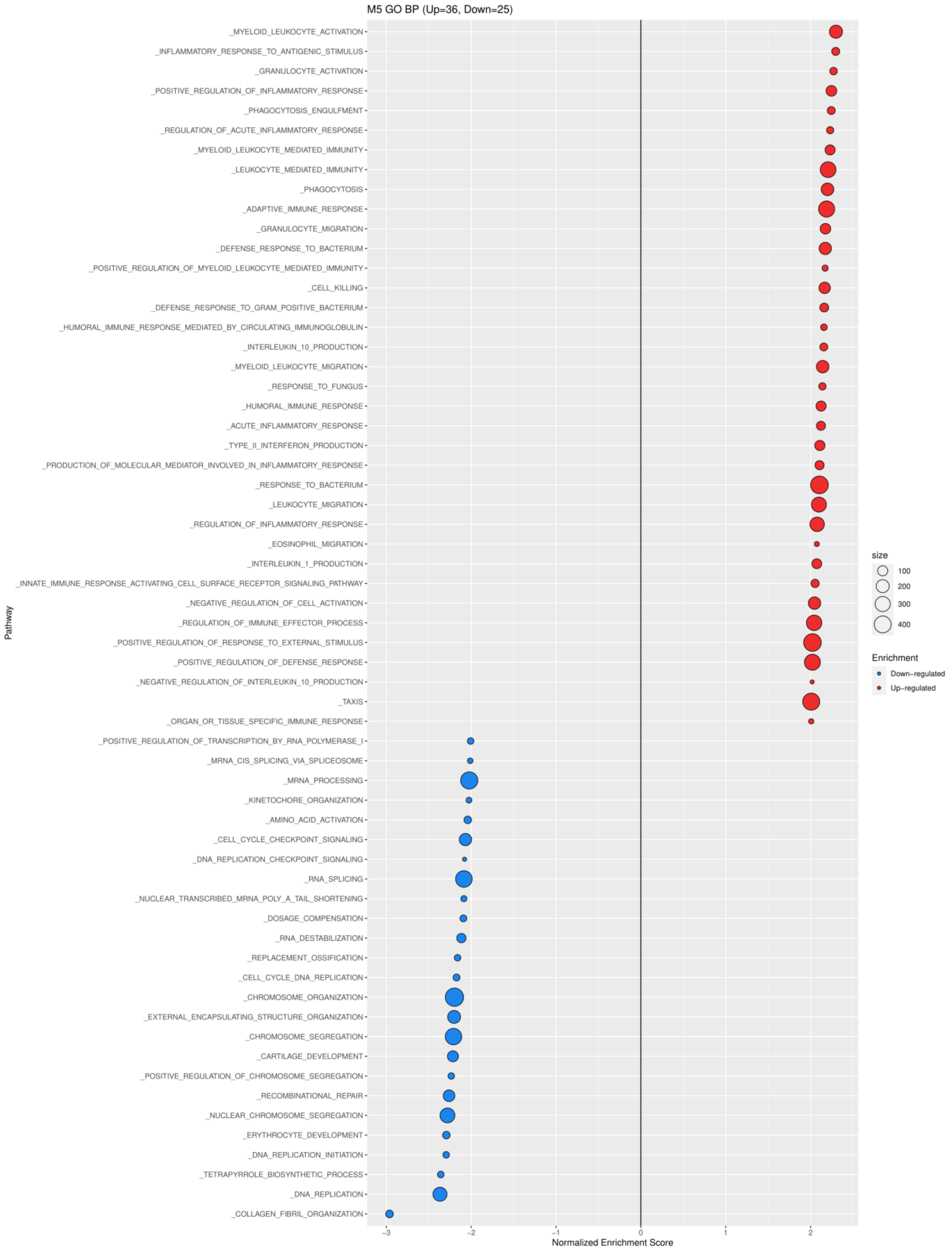


**Supplementary Figure 3:** M5 GO BP pathway analysis of SASH vs WT. N=5 mice/arm.

### Supplemental Figure 4: SASH BM CD45$^{-}$ stromal cells exhibit similar gene expression profiles to WT mice

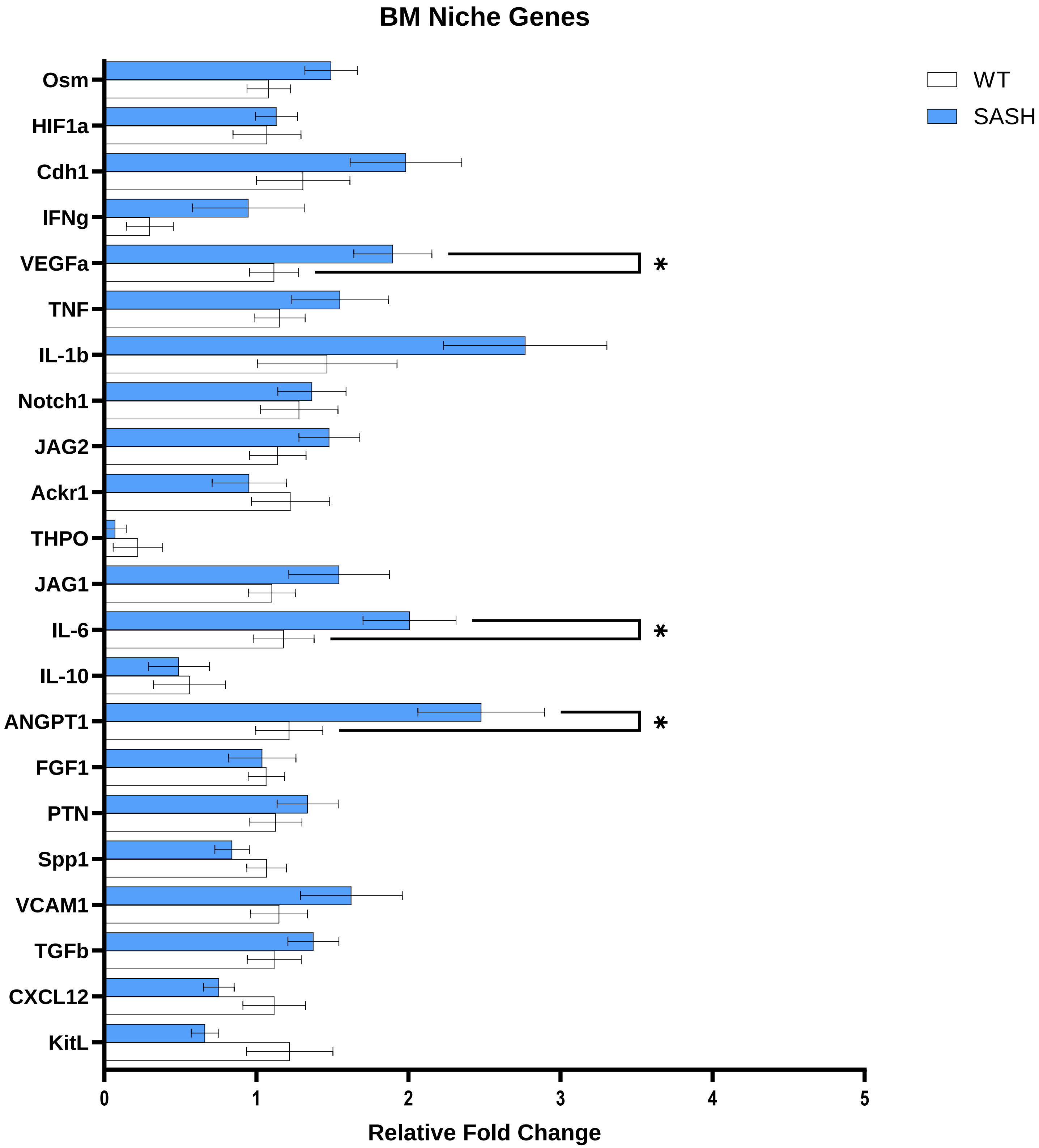


**Supplementary Figure 4:** RT-PCR analysis of niche factor genes within BM stromal cells. N=10 mice/arm. Error bars represent SEM. Students two-tailed T test was used for statistical analysis with p>0.05=*, p>0.01=**, p>0.001=***,p>0.0001=****

### Supplemental Figure 5: Bulk and stromal expression of HDC and histamine receptors in WT and SASH mice

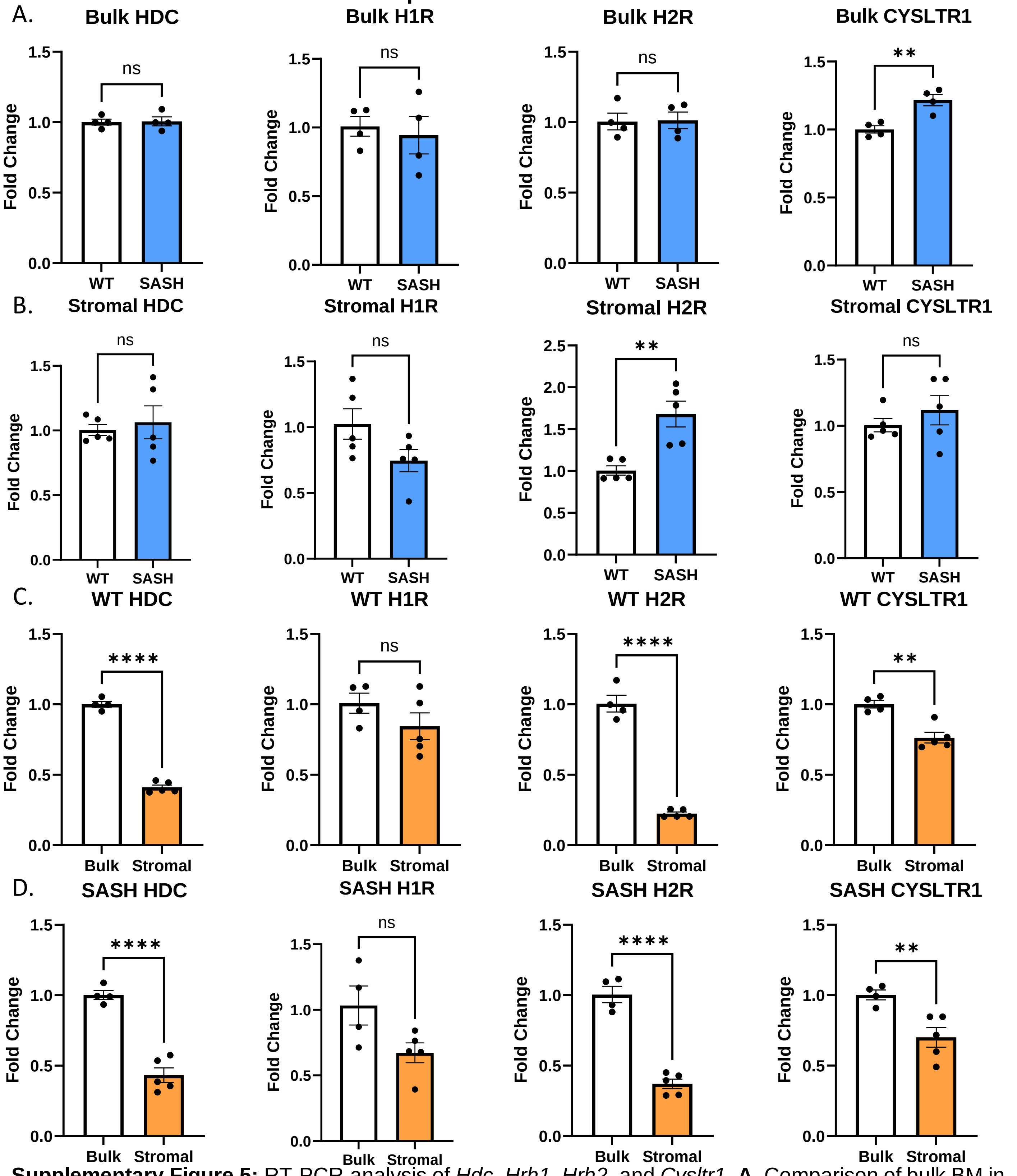


**Supplementary Figure 5:** RT-PCR analysis of *Hdc*, *Hrh1*, *Hrh2*, and *Cysltr1*. **A**. Comparison of bulk BM in WT and SASH mice. N=4 mice/group. **B.** Comparison of stromal BM in WT and SASH mice. N=5 mice/arm. **C.** Comparison of bulk and stromal BM in WT mice. N=4-5 mice/group. **D.** Comparison of bulk and stromal BM in SASH mice. N=4-5 mice/group. **A-D.** Error bars represent SEM. Students two-tailed T test was used for statistical analysis with p>0.05=*, p>0.01=**, p>0.001=***,p>0.0001=****

### Supplemental Figure 6: PB and BM kinetics of histamine administration in SASH mice

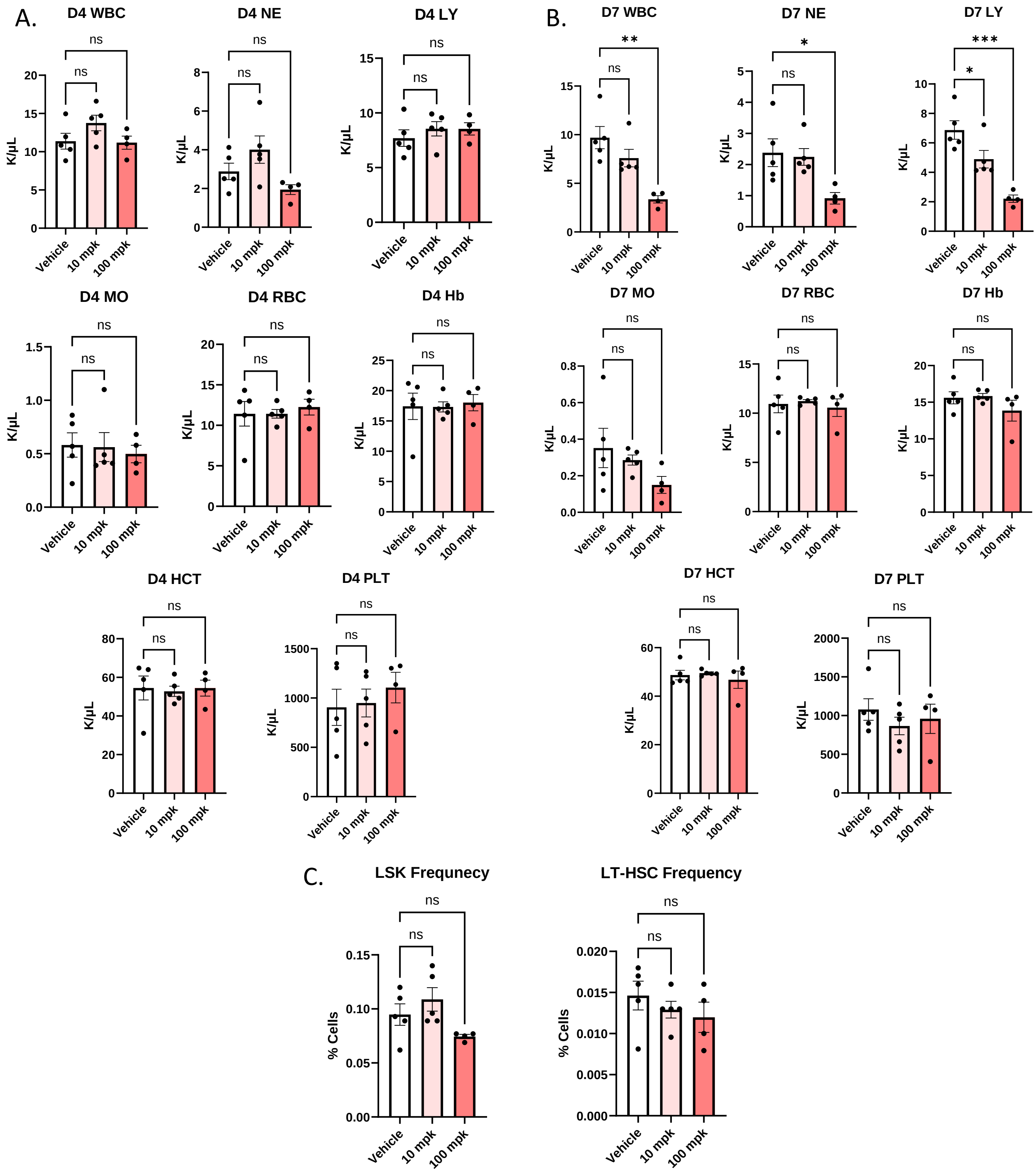


**Supplementary Figure 6:** Complete blood count analysis of circulating WBC, NE, LY, MO, RBC, Hb, HCT, and PLT after BID administration of 10 and 100 mg/kg histamine at **A.** D4 and **B.** D7. N=4-5 mice/group. **C.** Immunophenotypic analysis of HSPCs and LSKs per hindlimb after 7 days of BID 10 and 100 mpk histamine administration. N=4-5 mice/arm. **A-C.** Error bars represent SEM. One-way ANOVA was used for statistical analysis with p>0.05=*, p>0.01=**, p>0.001=***,p>0.0001=****

### Supplemental Figure 7: HSPC flow cytometry analysis gating strategy

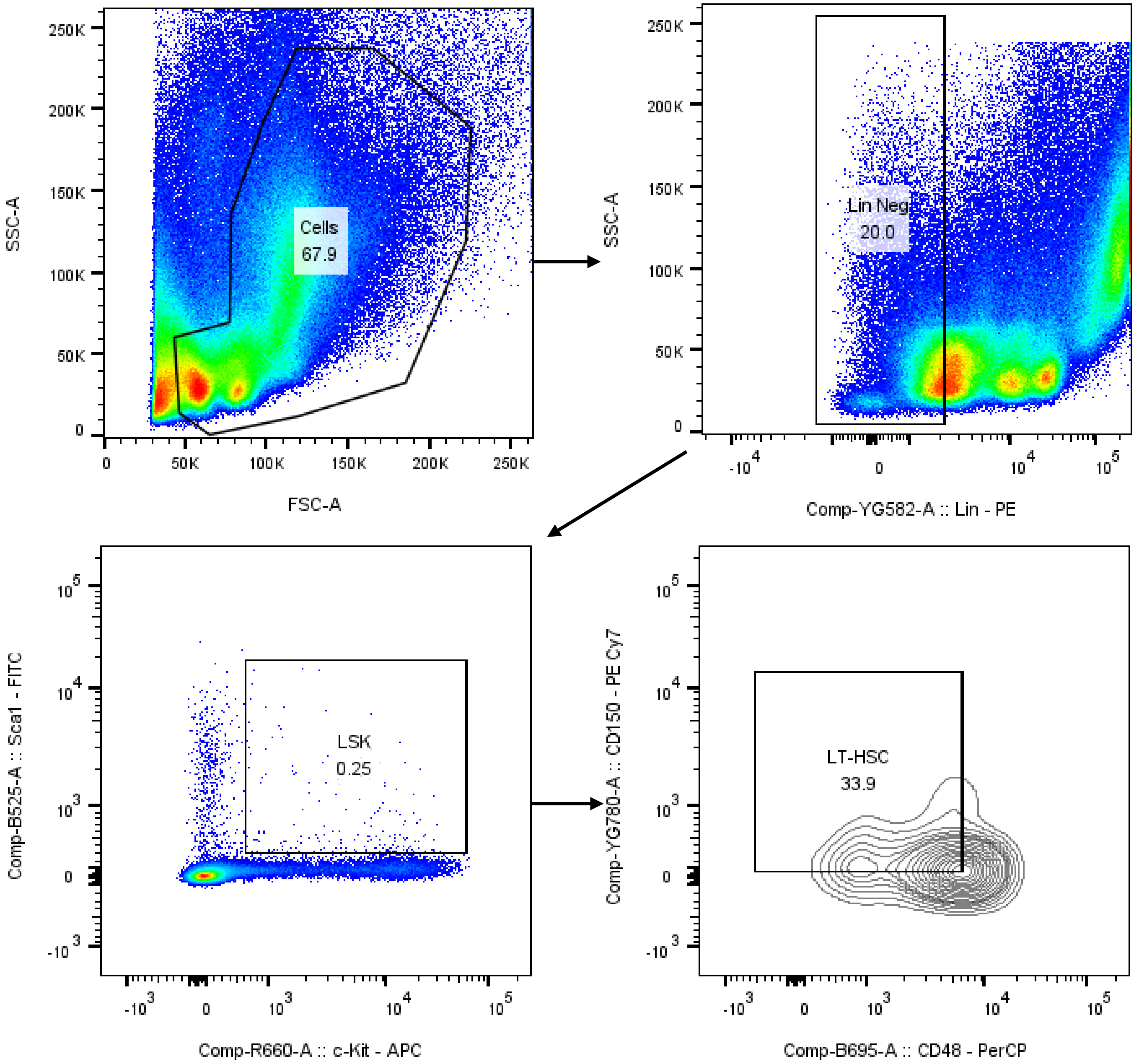


**Supplementary Figure 7:** Gating strategy for HSPC cell frequency analysis

### Supplemental Figure 8: Myeloid and lymphoid flow cytometry analysis gating strategy

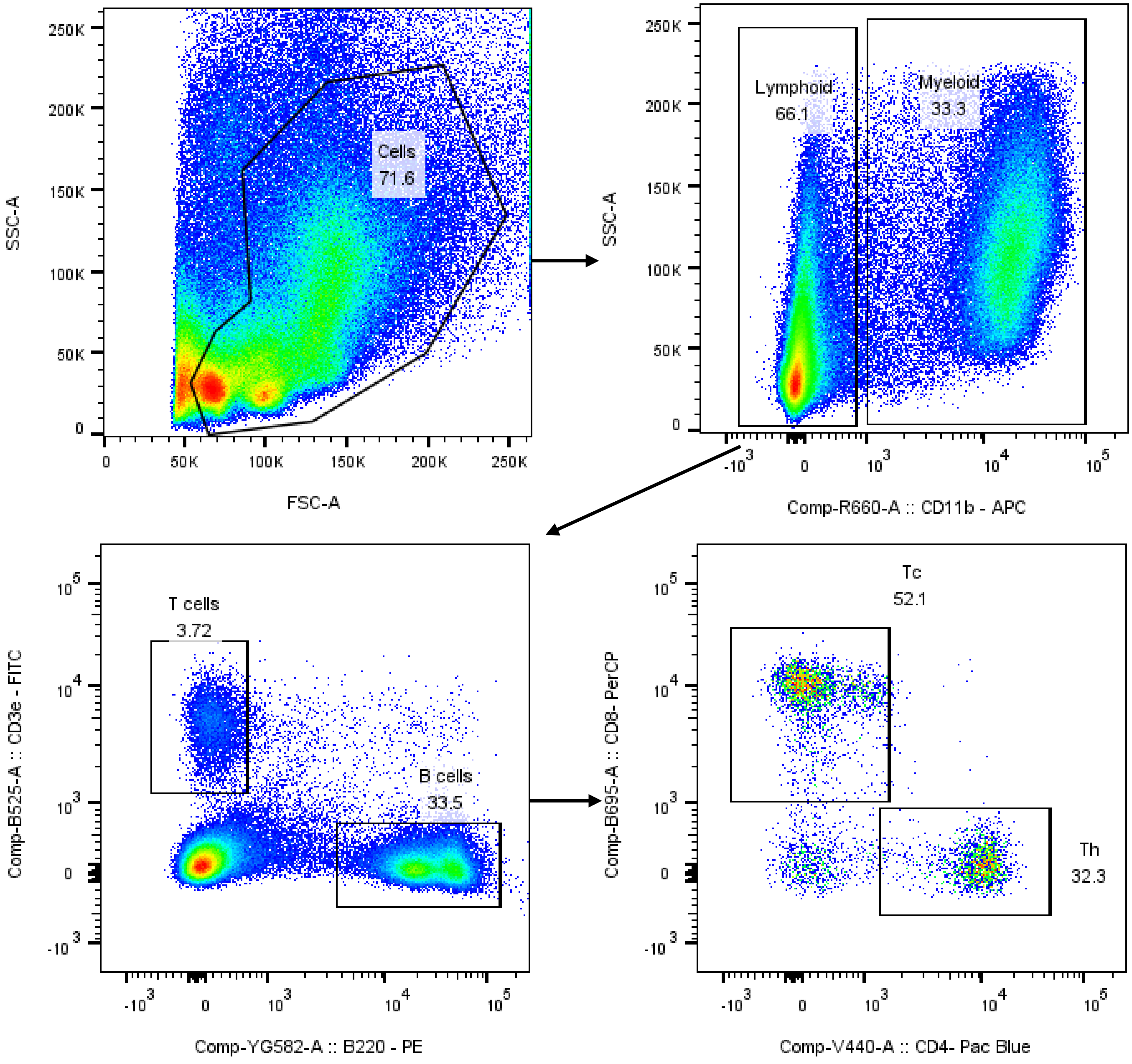


**Supplementary Figure 8:** Gating strategy for Myeloid and lymphoid cell frequency analysis.

### Supplemental Figure 9: CD45$^-$ cell flow cytometry analysis gating strategy

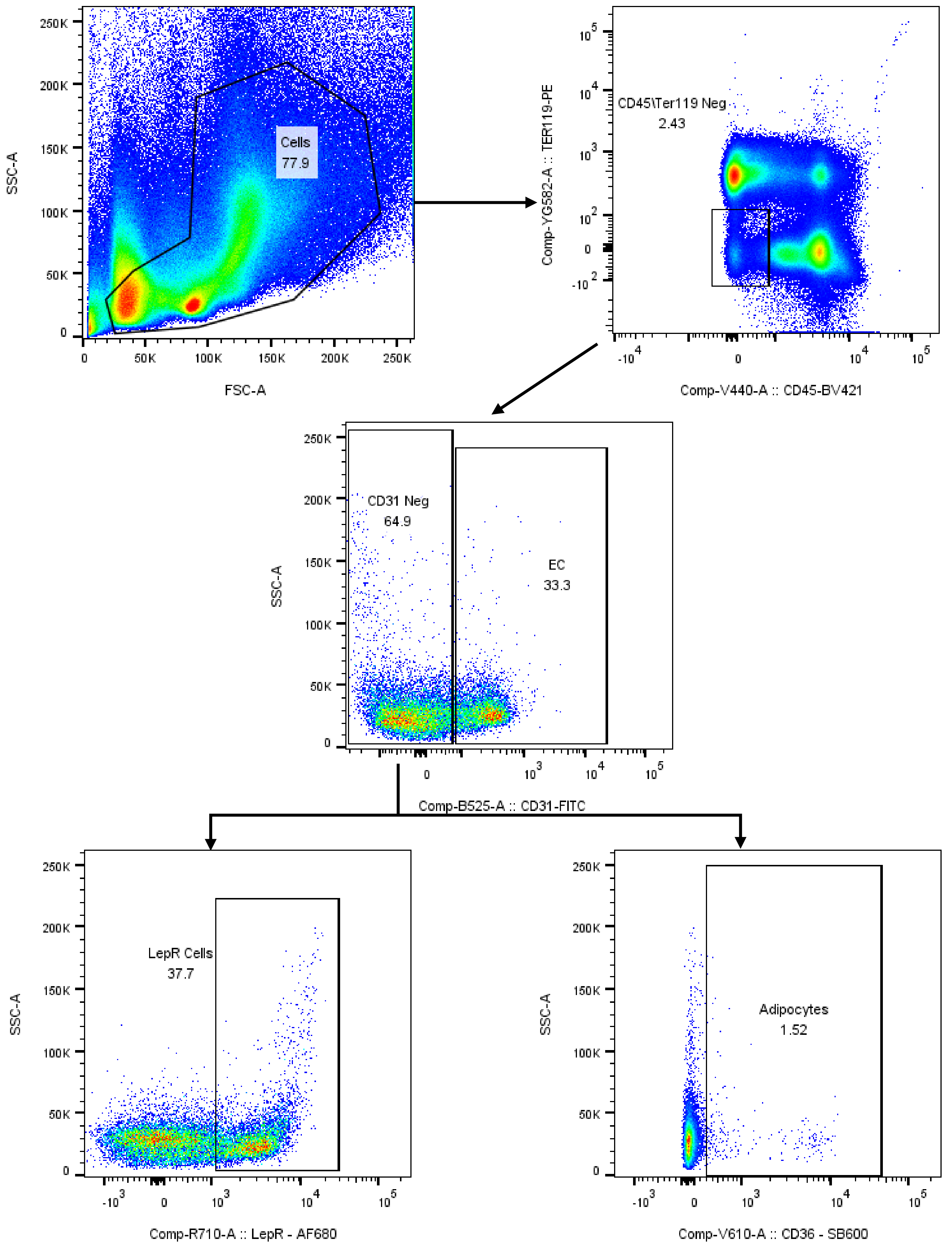


**Supplementary Figure 9:** Gating strategy for CD45$^-$ cell frequency analysis.

#### Supplemental Methods

**Flow Cytometry:** Cells were stained for myeloid and lymphoid populations with antibodies against CD11b (M1/70), CD45R/B220 (RA3-6B2), CD3e (17A2), CD4 (RM4-4), and CD8a (53-6.7). Cells were stained for stem and progenitor populations with antibodies against CD11b (M1/70), CD45R/B220 (RA3-6B2), CD3e (17A2), Ly-6G and Ly-6C (RB6-8C5), TER-119 (TER-119), CD117 (2B8), Ly-6A/E (D7), CD48 (HM48-1), and CD150 (mShad150). Cells were stained for $CD45^-$ populations with antibodies against CD45 (30-F11), TER-119 (TER-119), CD31 (390), S100A4 (2G11B4), CD146 (P1H12), CD36 (HM36), CD51 (RMV-7), CD140a (APA5), and Leptin Receptor (JA73-01).

**RNA Isolation and Quantitative PCR:** Cell separations were performed using mouse anti-antibody microbeads (Miltenyi Biotec) and separated into specific cell lineages by magnetic bead separation per manufacturer's protocol. RNA was isolated using the RNeasy Plus Mini Kit (QIAGEN cat # 74136) according to the manufacturer's protocol. Isolated RNA concentration and integrity was evaluated with a TapeStation Instrument (Agilent). RNA was then converted to cDNA using Takara primescript RT reagent kit (VWR cat # 102940-202) following the manufacturer's instructions. Real-time PCR measurement was performed in a 20 µL reaction containing 1 µL cDNA template in a 1:20 dilution of primer/probe with 1X Accuris Taq DNA polymerase. Samples were run on a CFX96 optical module (Bio-Rad). Thermal cycling conditions were 95C for 3 minutes, followed by 50 cycles of 95C for 15 seconds and 60C for 1 minute. Murine probe/primer sets for all genes assayed were obtained from Life Technologies. For each reverse transcription reaction, Cq values were determined as the average values obtained from three independent real-time PCR reactions.

**RNA Sequencing:** RNA samples were shipped on dry ice to Azenta for subsequent library preparation and PE150 sequencing on Illumina (35M paired-end reads/sample). Meticulous implemented quality control was assessed covering per base quality score, reads quality score, and reads GC distributions. Subsequently, sequence reads underwent trimming to eliminate potential adapter sequences and nucleotides with poor quality, utilizing Trimmomatic v.0.36[1]. Following this preprocessing step, alignment of the reads to the reference genome (Mus musculus GRCm38 from ENSEMBL) was carried out using the STAR aligner[2], and unique gene hit counts were derived using featurecounts[3]. The resulting gene hit counts data underwent normalization for Principal Component Analysis (PCA). DESeq2[4] was then employed to compare gene expression across different sample groups, with the Wald test generating p-values and log2 fold changes. Genes meeting the criteria of an adjusted p-value $< 0.05$ and an absolute log2 fold change $> 1$ were deemed as differentially expressed genes for each comparison. Furthermore, the 'complexheatmap' tool in R[5] was employed to produce various heatmaps. Finally, Gene Set Enrichment Analysis (GSEA)[6] was utilized to establish a priori defined sets of genes exhibiting statistically significant, concordant differences between two biological states, leveraging the Molecular Signatures Database (MSigDB)[7].

**TriNetX Patient EHR Analysis:** TriNetX is a federated health research network that provides access to electronic health records (EHRs) from approximately 100 million unique patients across the United States and has been utilized in an array of large-scale clinical research studies. The platform provides web-based secure access to patient EHR data from hospitals, primary care clinics, and specialty treatment providers. Available data includes demographics, diagnoses, procedures, medications, laboratory tests, vital signs, and healthcare utilization. TriNetX reports population-level data without revealing protected health information; consequently, the MetroHealth System, Cleveland Ohio, Institutional Review Board (IRB) determined that research using TriNetX is not Human Subject Research and IRB review is not required. All patients included in the analysis had a diagnosis of allergy (based on International Classification of Diseases Tenth Revision or ICD-10) code T78) who were subsequently prescribed antihistamines or other anti-allergy medications (decongestants and antiallergics, corticosteroids). We investigated the rate of the diagnosis of elevated white blood cell counts in patients prescribed antihistamines vs other anti-allergy medications. Drug exposure information is recorded as RxNorm codes in TriNetX. The outcome of interest is the diagnosis of elevated white blood cell count (D72.82) within one year of drug prescription. The study population was divided into two cohorts. The exposure cohort was comprised of patients with no history of elevated white blood cell count who were prescribed the antihistamines following the diagnosis of allergy. The control cohort included patients with no history of elevated white blood cell count who were prescribed other anti-allergy medications following a diagnosis of allergy. The index event for each candidate drug evaluation was the initial date of drug prescription.

The exposure and control groups were propensity-score matched for baseline covariates including demographics (age, gender, race/ethnicity), adverse determinants of socioeconomic factors, and comorbidities including hematologic cancers and other medications. The TriNetX built-in propensity score matching function was used, which involves 1:1 matching using a nearest neighbor greedy matching algorithm with a caliper of 0.1 standardized mean differences (SMD) to account for potential confounding variables. The outcome "elevated white blood cell counts" that occurred within a 12-month follow-up period were compared between the matched exposure and control cohorts using Cox proportional hazard analyses with censoring applied. Hazard ratios (HR) and 95% confidence intervals (CI) were calculated. All the analyses were performed within the TriNetX Analytics Platform using built-in functions.
